# Advancing Understanding of DNA-BfiI Restriction Endonuclease Cis and Trans Interactions through smFRET Technology

**DOI:** 10.1101/2023.03.31.535070

**Authors:** Šarūnė Ivanovaitė, Justė Paksaitė, Aurimas Kopūstas, Giedrė Karzaitė, Danielis Rutkauskas, Arunas Silanskas, Giedrius Sasnauskas, Mindaugas Zaremba, Stephen K. Jones, Marijonas Tutkus

## Abstract

Monitoring of DNA-protein interactions is essential in understanding many biological processes. Proteins must find their target site on a DNA molecule to perform their function, and the mechanisms for target search differ across proteins. Revealing temporal interactions with two target sites, both in *Cis* and in *Trans*, is crucial in target search mechanisms studies. Here, we present two single-molecule Förster resonance energy transfer (smFRET)-based assays to study BfiI-DNA interactions. The first assay, smFRET-based DNA looping assay, detects both “Phi” and “U”-shaped DNA looping events. We modified it to only allow *in Trans* BfiI-target DNA interactions to improve specificity and reduce limitations in the observation time. Our TIRF microscopy measurements directly observe the on- and off-target binding events and characterize BfiI binding events. Our results show that BfiI binding events last longer on target sites and that the BfiI rarely changes conformations during binding. This newly developed assay could be useful for other two-targets-binding DNA-interacting proteins and could be employed for dsDNA substrate BfiI-PAINT, which is useful for DNA stretch-assays and other super-resolution fluorescence microscopy studies.

## Introduction

DNA-protein interactions play a key role in many biological processes^1,2^ and occur at nanometer lengths and millisecond-to-second timescales. One aspect unifying these interactions is that a protein must find its target site on a DNA molecule to perform its function: cleavage, modification, activation of other proteins, etc.^3^ Target search mechanisms differ across proteins, but can be divided into several classes: 3D diffusion (i.e., jumping), sliding, hopping, and intersegmental transfer.^4,5^ They also differ based on the number of DNA target sites that a protein binds. Proteins with a single DNA target binding domain utilize one or combination of aforementioned target search mechanisms.^6–8^ However, proteins with two DNA-binding domains may simultaneously interact with two targets on a single DNA molecule (i.e. *in Cis* interaction) or on two separate molecules (i.e. *in Trans* interaction).^9^ Temporal interactions with two target sites *in Cis* and *in Trans* often play an important role in target search mechanisms and are detrimental for various cellular mechanisms^10^.

Continuous real-time monitoring of these dynamic interactions typically requires immobilizing one of interacting partners on a surface, after which they can be visualized in great detail through force spectroscopy, fluorescence, Förster resonance energy transfer (FRET), atomic force microscopy (AFM), nanopores, and DNA flow-stretch assays.^11^ Single molecule FRET (smFRET) approaches also work well for high-throughput format studies of many protein-DNA interactions^12^ including CRISPR-Cas ^13,14^, SSB protein ^15^, RecA^16^, DNA polymerase^17,18^, RNA polymerase^19,20^ and other proteins ^21–23^. The smFRET experiments can follow different study designs including: A) both dyes of a FRET pair on a single DNA fragment,^24^ B) one dye on DNA and the other on protein,^8^ and C) both dyes on a single protein.^25^ Notably, these designs are mainly employed to study DNA-protein interactions occurring in-cis, i.e. when both target sites are on a single surface-immobilized DNA molecule, but not for in-trans interactions.

We have previously described an smFRET approach, utilizing DNA-looping for studying DNA binding by Type IIF restriction endonucleases Ecl18kI and NgoMIV.^27^ Two Ecl18kI dimers bind two pseudo-palindromic DNA targets (5′-CCNGG-3′, N is any nucleotide) *in Cis*, which leads to protein tetramerization and formation of either of two possible DNA loop types - “U”-or “Phi”-shaped (Fig. 1A).^27^ In the “Phi”-shaped DNA loop, conformation distance between FRET pair fluorophores was too high for probing by FRET methods and like non-looped DNA, resulted in “zero” FRET efficiency. By simultaneously monitoring tethered fluorophore motion and FRET efficiency, we revealed a “Phi”-shaped DNA loop and true non-looped DNA states.^26^ The homotetrameric REase NgoMIV^29,30^ can simultaneously bind two palindromic (5’-GCCGGC-3’) target sites and can form both types of loops (Fig. 1B).^28^ In our study, both DNA loop shapes resulted in non-zero FRET efficiency. However, NgoMIV limits our assay, since increased protein concentration increases the probability of having two tetramers per single DNA fragment containing two target sites. This, in turn, limits DNA looping and introduces nonspecific binding events that are more difficult to interpret.

**Figure 1:**
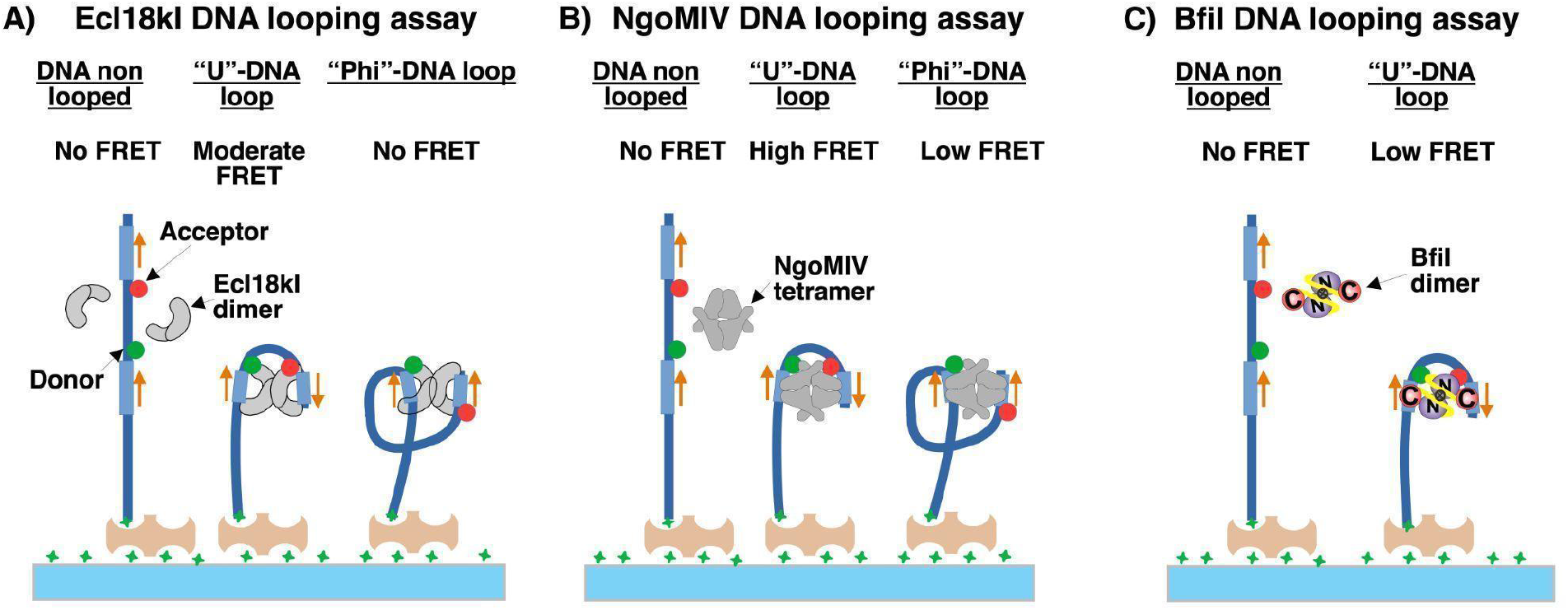
Schematic diagram illustrating a single-molecule FRET-based DNA looping assay for probing *in Cis* DNA-protein interactions across different restriction endonucleases. **A)** Ecl18kI and tetramer-DNA interact and form “U”- and ‘‘Phi’’-shaped DNA loops. Signals corresponding to different DNA conformations are depicted.^26,27^ **B)** NgoMIV and homotetramer-DNA interact and form ‘‘U’’- and ‘‘Phi’’-shaped DNA loops.^28^ **C)** BfiI and dimer-DNA interact and form a “U”-shaped DNA loop.

In order to further expand our smFRET-based assay for studying protein-DNA interactions, we conducted studies of the type IIS REase BfiI. In contrast to Ecl18kI and NgoMIV, the BfiI homodimer contains separate domains for cleavage and recognition (Fig. 2A). Binding of cognate DNA to BfiI presumably triggers a conformational change from the “closed” to “open” state and opens an entrance to the active site in the N-terminal domains.^31–34^ BfiI REase requires two copies of an asymmetric 5’-ACTGGG-3’ target site,^35,36^ and therefore it can only make one type of DNA loop (Fig. 1C, Fig. 2B). BfiI does not require Mg^2+^ ions for catalysis,^37^ therefore we chose to study BifI protein with a K107A active site mutation ^36^ (hereinafter in this text called BfiI protein). Here we employed two different smFRET schemes to measure BfiI-DNA interactions. By directly observing on-/off-target binding events *in Cis* and *in Trans* we were able to characterize their durations and frequency. In addition, we described and demonstrated a novel smFRET based approach to study protein-DNA interactions *in Trans* which could prove useful for studying other DNA binding proteins in the future.

**Figure 2:**
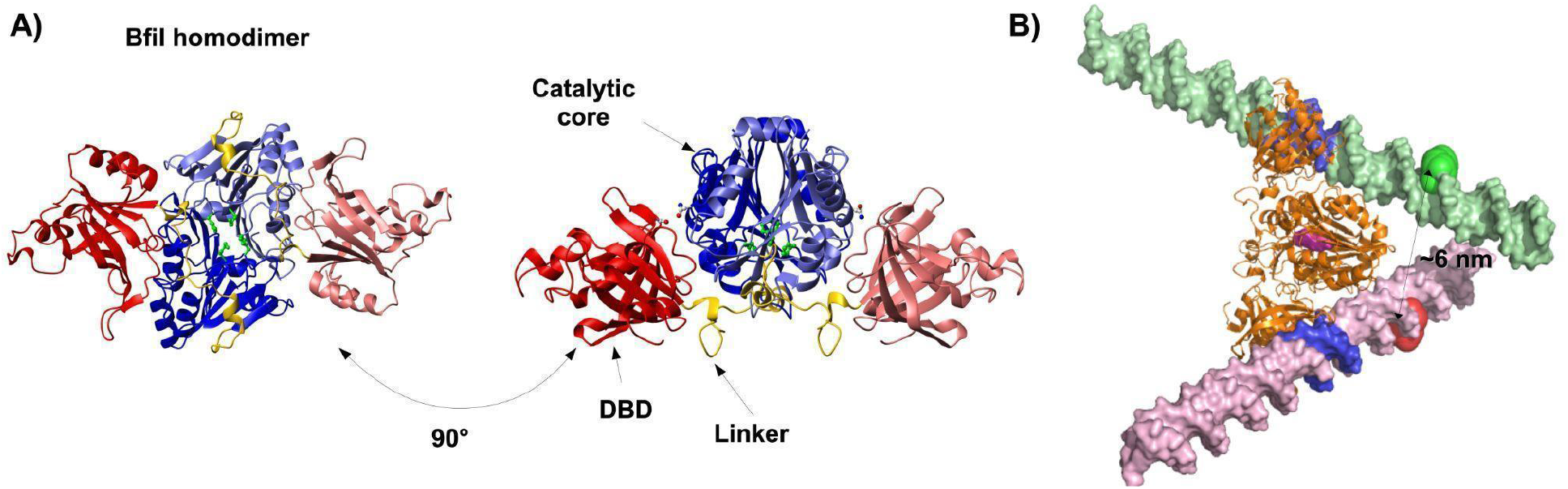
Overall BfiI structure without and with DNA substrates. **A)** BfiI homodimer crystal structure shown in two different orientations (PDB ID: 2C1L). Each monomer is composed of the N-terminal PLD-like domain and the C-terminal DNA binding domain (DBD). Two N-terminal domains (light and dark blue) make a dimeric catalytic core flanked by two C-terminal DBDs (light and dark red). The linker connecting the N- and C-terminal domains is colored yellow. The active site residues are shown in green-colored balls–and-sticks. The K107A mutation makes BfiI catalytically inactive. **B)** Model of BfiI homodimer in complex with two double-stranded DNA molecules containing target sites based on the co-crystal structure of BfiI DBD with cognate DNA (PDB ID: 2C1L).^32^ Target sites on DNA molecules are marked in blue, position of donor fluorophore on DNA is marked by green-colored spheres, and position of acceptor fluorophore on DNA is marked by red-colored spheres. The active site of BfiI is marked by magenta-colored spheres.

## Materials and methods

### Protein and DNA fragments

The active site mutant BfiI was purified as described previously.^36^ Protein concentrations are expressed in terms of dimer. We synthesized 350 bp DNA fragments containing either one (for *in Trans* interaction studies) or two target (for *in Cis* interaction) sites for BfiI: 5’-ACTGGG-3’. The scheme for fragment synthesis was the same as described previously.^26^ First, auxiliary DNA fragment for 150 bp inter-target distance was produced using pUC19 as a template and 5’-ACTGGGCTGTCTATTATTATTGCAGCAGCCACTGGTAAC-3’, 5’-TGGGCTGCATTTATTATTATTGAGCGCAGATACCAAATAC-3’ as primers. A biotinylated fragment was produced from pUC19 template using 5’-TAGTGCACGCGGTGTTCGCTCCAAGCTGGGCTGTG-3’ and 5’-CGTATGTCGTACCGGTAAGAACTCTGTAGCACCGCC-3’primers.

DNA fragments for *in Cis* interaction studies (i.e. DNA looping): FRET pair-labeled fragment was produced using an auxiliary DNA fragment as a template and the following two primers 5’-AGCGTAGC**ACTGGG**CTGTCTATTA[ATTO647N]TATTGC-3’, 5’-GAGCACCGCGTGTAGC**ACTGGG**CTGCATTTAT[Cy3B]ATTATTG-3’. Next, the biotinylated and FRET pair-labeled fragments were digested by AdeI, ligated, purified and stored at -20 °C until use.

DNA fragments for *in Trans* interaction studies: FRET pair-labeled single BfiI target site containing DNA fragment was produced using using an auxiliary fragment as a template and the following two primers: 5’-AGCGTAGC**ACTGGG**CTGTCTATTA[ATTO647N]TATTGC-3’, 5’-GAGCACCGCGTGTAGCCCTGGGCTGCATTTAT[Cy3B]ATTATTG-3’. Next, the biotinylated and FRET pair-labeled fragments were AdeI digested, ligated, purified and stored at -20 °C until use.

The Cy3B-labeled dsDNA oligonucleotide (Cy3B-oligo) containing BfiI target site was assembled by annealing oligonucleotides with these sequences: 5’-GAGCACCGCGTGTAGC**ACTGGG**CTGCATTTAT[Cy3B]ATTATTG-3’and 5’-CAATAATAATAAATGCAG**CCCAGT**GCTACACGCGGTGCTC-3’.

The Cy3B-labeled dsDNA oligonucleotide without the BfiI target site was assembled by annealing oligonucleotides with these sequences:

5’-GAGCACCGCGTGTAGCCCTGGGCTGCATTTAT[Cy3B]ATTATTG-3’and 5’-CAATAATAATAAATGCAG**CCCAGT**GCTACACGCGGTGCTC-3’

### Sample preparation

The flowcell was assembled from a six-channel Sticky-Slide VI 0.4 (Ibidi, Germany) and a PEG-coated coverslip (Menzel Glaser, Braunschweig, Germany). The procedure of the coverslip modification with PEG derivatives was described previously.^38^ The flowcell channel was incubated with 0.5 mg/ml of neutravidin (A-26666, Molecular Probes) in yellow buffer (YB) (33 mM Tris-acetate (pH 7.9 at 20°C), 66 mM K-acetate, 1.5 mM BSA) for 3 min, washed with YB, incubated with ∼0.1 nM DNA in YB for 10 min, and washed with YB. For the measurement of the DNA-protein interactions, the cell was infused with 0.5 nM BfiI and 0.2 nM of Cy3B-labeled dsDNA oligonucleotide in imaging buffer (IB, the YB supplemented with 15 units/mL of glucose oxidase (G6125, Sigma-Aldrich), 1% of glucose (G0047, TCI Europe), 120 units/mL of catalase (C9322, Sigma-Aldrich), and 2.5 mM of UV-treated Trolox (Tx, 238813, Sigma-Aldrich). Each different condition reported in this work was measured in a separate channel of the flowcell.

### Single-molecule data acquisition

An objective-type TIRF microscope was used for single-molecule fluorescence movie acquisition.^38,39^ 532 and 635 nm laser powers were set to ∼2 mW after the objective and a ZT532/635rpc-XT dichroic mirror (Chroma) was used in the filter turret of the microscope. The fluorescence was filtered with a quadruple-band interference filter FF01-446/510/581/ 703 (Semrock), split by T640lpxr-UF2 (Chroma), and imaged with an EMCCD (DU-897E-CS0-UVB, Andor) with 100 ms integration time.

### Data analysis

In microscopy movies, fluorescent molecules were identified, and intensity versus time trajectories were extracted using the custom analysis package written in Igor Pro (Wavemetrics, Portland, OR) as previously described.^26,40^ Time trajectories were screened to exhibit characteristic single-molecule FRET features: clear donor-acceptor anti-correlated intensity changes, a typical single-molecule fluorescence intensity, and a single bleaching step. Protein binding events were detected in the intensity versus time trajectories automatically and then examined manually according to these criteria: intensity of donor at 532 nm excitation should be higher than 30 a.u., the binding event should last for longer than 5 frames.

## Results and discussion

### The SM assay for BfiI-DNA *in Cis* interaction

For BfiI protein and DNA *in Cis* interaction studies, we employed our smFRET-based DNA looping assay.^26^ The first scheme monitors *in Cis* interactions and FRET efficiency changes within two target sites on a single surface-immobilized DNA fragment upon BfiI-induced DNA looping. First, we immobilized biotinylated dsDNA molecules containing two BfiI target sites on a PEGylated glass coverslip surface via neutravidin. The first target site was labeled with a Cy3B fluorophore, and the second target site with an ATTO647N fluorophore. In the presence of BfiI protein, we expected to observe two DNA states: looped and non-looped. The non-looped DNA state should result in lowest possible apparent FRET efficiency, which we termed the “zero” FRET level. Since BfiI can bind its non-palindromic recognition sites in one orientation only, the DNA should adopt a “U”-shaped conformation upon protein-induced looping and the FRET efficiency should increase. After binding both target sites, the BfiI protein may undergo a conformational transition from “closed” to “open”. Upon such conformational change we should observe a decrease in FRET efficiency. However, it is very likely that the “open” conformation and the non-looped DNA would both have “zero” FRET level, since modeling data predicts that the fluorophores on DNA when it is bound by BfiI would be separated by > 10 nm. Also, when BfiI is bound to a single target site, we would observe no FRET efficiency increase compared to protein-free DNA.

To measure the value of apparent FRET efficiency in non-looped state, we acquired a series of images of the DNA sample immobilized on the flowcell surface without and with the BfiI protein using a home-built TIRF set-up (see Materials and methods) and 532 nm laser excitation. Most observed signals with and without BfiI showed characteristic SM behavior: strong donor signal intensity, and single-step bleaching events (Fig. 3B). FRET efficiencies from the extracted SM signals without BfiI revealed a single Gaussian FRET efficiency distribution centered at ∼0.1 FRET efficiency. In the presence of BfiI, the FRET efficiency distribution had an overlapping double-Gaussian shape. The first Gaussian we attributed to the non-looped DNA state because it coincided with the protein-free control (both centered at ∼0.1 FRET efficiency). Since the first Gaussian coincided with the protein-free control (both centered at ∼0.1 FRET efficiency), we attributed it to the non-looped DNA state. The second Gaussian had ∼0.08 higher FRET efficiency, indicating the looped DNA state (Fig. 3C).

**Figure 3:**
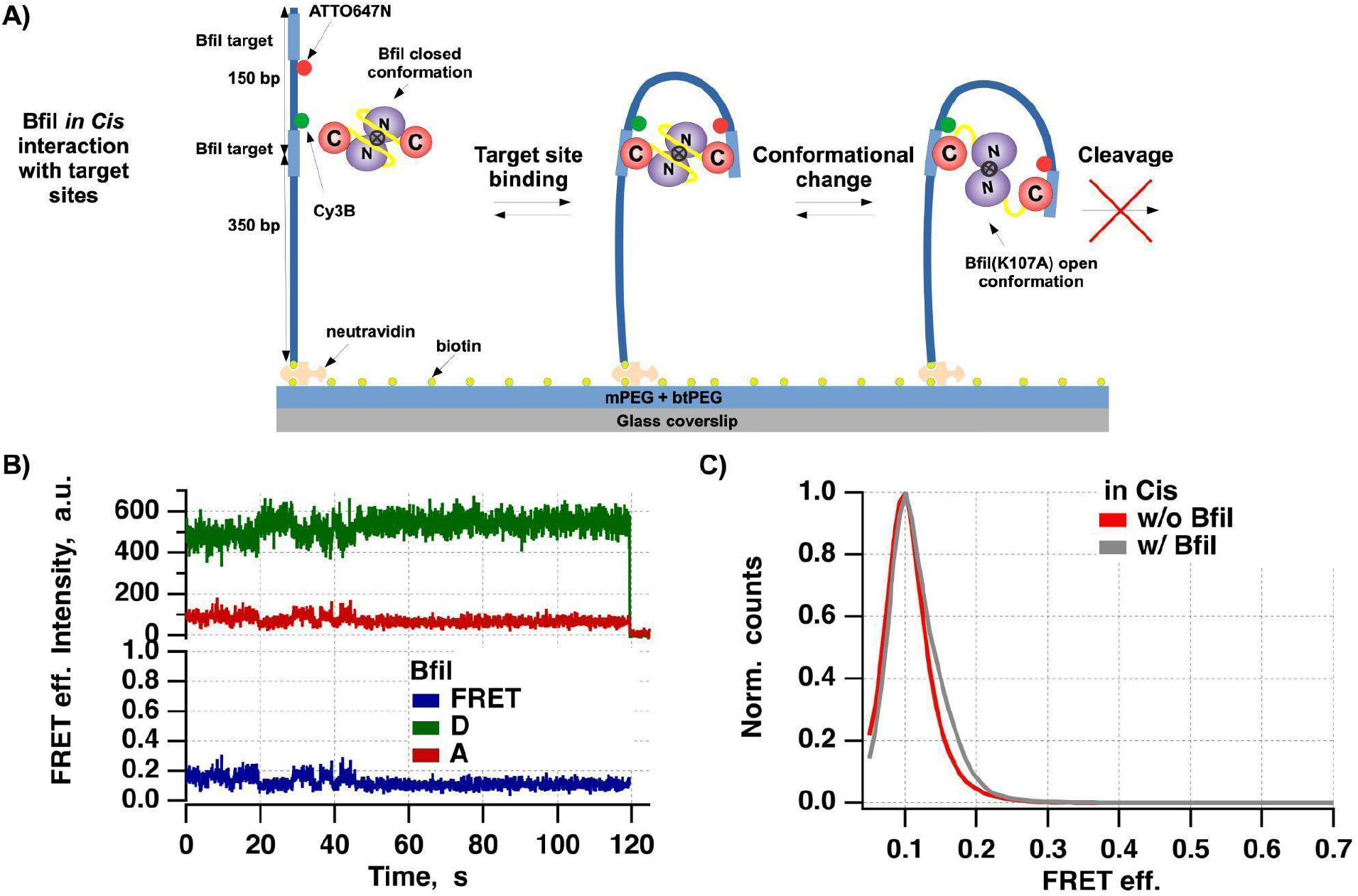
Results of single-molecule FRET (smFRET) assay for BfiI-DNA in Cis interaction studies. **A)** Diagram illustrating the scheme of smFRET-based DNA looping assay. Biotinylated double-stranded DNA fragments containing two BfiI target sites were immobilized on PEGylated (10% biotin-PEG) surface via neutravidin. The target site located closer to the surface has Cy3B fluorophore, and the surface distant site contains ATTO647N. Binding of BfiI protein to both target sites induces DNA looping and brings these two fluorophores in a close proximity, and increases FRET efficiency. Subsequent conformational changes of BfiI between closed to open states, may reduce the observed FRET efficiency. Active site mutation of BfiI makes it catalytically inactive and prevents target site cleavage. **B)** Representative SM signals: donor intensity (green), acceptor intensity (red), and apparent FRET efficiency (blue), showing DNA-BfiI interaction in Cis. **C)** Graph showing a distributions of FRET efficiencies of selected SM signals with (w/) and without (w/o) BfiI protein.

As predicted, we saw an increase of FRET efficiency in the presence of BfiI, which we interpreted as DNA loop formation due to binding of both targets. Therefore, in this assay “zero” FRET events could reflect several fundamentally different situations: 1) no protein bound to DNA, 2) single target site-bound BfiI, 3) two-target site-bound BfiI in the “open” conformation, 4) various off-target site BfiI-DNA interactions. While, the non “zero” FRET events could reflect any of BfiI conformation. Due to the ambiguity of the “zero” FRET events we mentioned above, we could not outrule the presence of other experimental states. Elimination of said ambiguity required a different experimental approach.

### The SM assay for BfiI-DNA interaction *in Trans*

To reveal BfiI-DNA interactions occurring only on the target site of surface-immobilized DNA molecules and to eliminate the possibility of DNA looping, we restricted our assay to allow *in Trans* interactions only by separating the asymmetric BfiI target between two dsDNA molecules. For this assay we employed surface immobilized biotinylated dsDNA molecules containing a single BfiI target site and labeled with ATTO647N (near the target site at the surface-distant end of the fragment) and Cy3B (near the middle of the DNA molecule). Co-localization of these two dyes allowed us to locate surface-immobilized dsDNA molecules more reliably than with a single fluorophore. Alongside the double labeled dsDNA, we used short Cy3B labeled dsDNA oligonucleotides (Cy3B-oligo), which contained the second BfiI target site. These short Cy3B labeled oligonucleotides were preincubated in a test tube with BfiI prior to experiments to form BfiI-Cy3B-oligo complexes.

We hypothesized that this modified assay would exclude the possibility of DNA looping and enable monitoring of binding events for longer periods with less problematic Cy3B bleaching. In our previously developed smFRET-based DNA looping assay donor and acceptor fluorophores were always present, therefore bleaching of the donor limited the meaningful observation time. Whereas here, each new BfiI binding event on surface-immobilized DNA would likely occur with a different Cy3B fluorophore, allowing us to monitor the protein-DNA interactions for longer periods.

In principle, BfiI can bind either on-or off-target on the surface-immobilized DNA molecules. We further expected that in this assay on-target binding of BfiI-Cy3B-oligonucleotide complex would give a higher than “zero” FRET efficiency and moderate Cy3B fluorescence intensity (Fig. 4A). In contrast, its binding to any off-target location of the immobilized DNA should result in high Cy3B fluorescence intensity and “zero” FRET efficiency. Closed to open state conformational transitions of BfiI should result in a shift from high to low or “zero” FRET efficiency. However, conformational changes of BfiI, occurring at off-target binding events, would not be manifested as changes in FRET efficiency.

**Figure 4:**
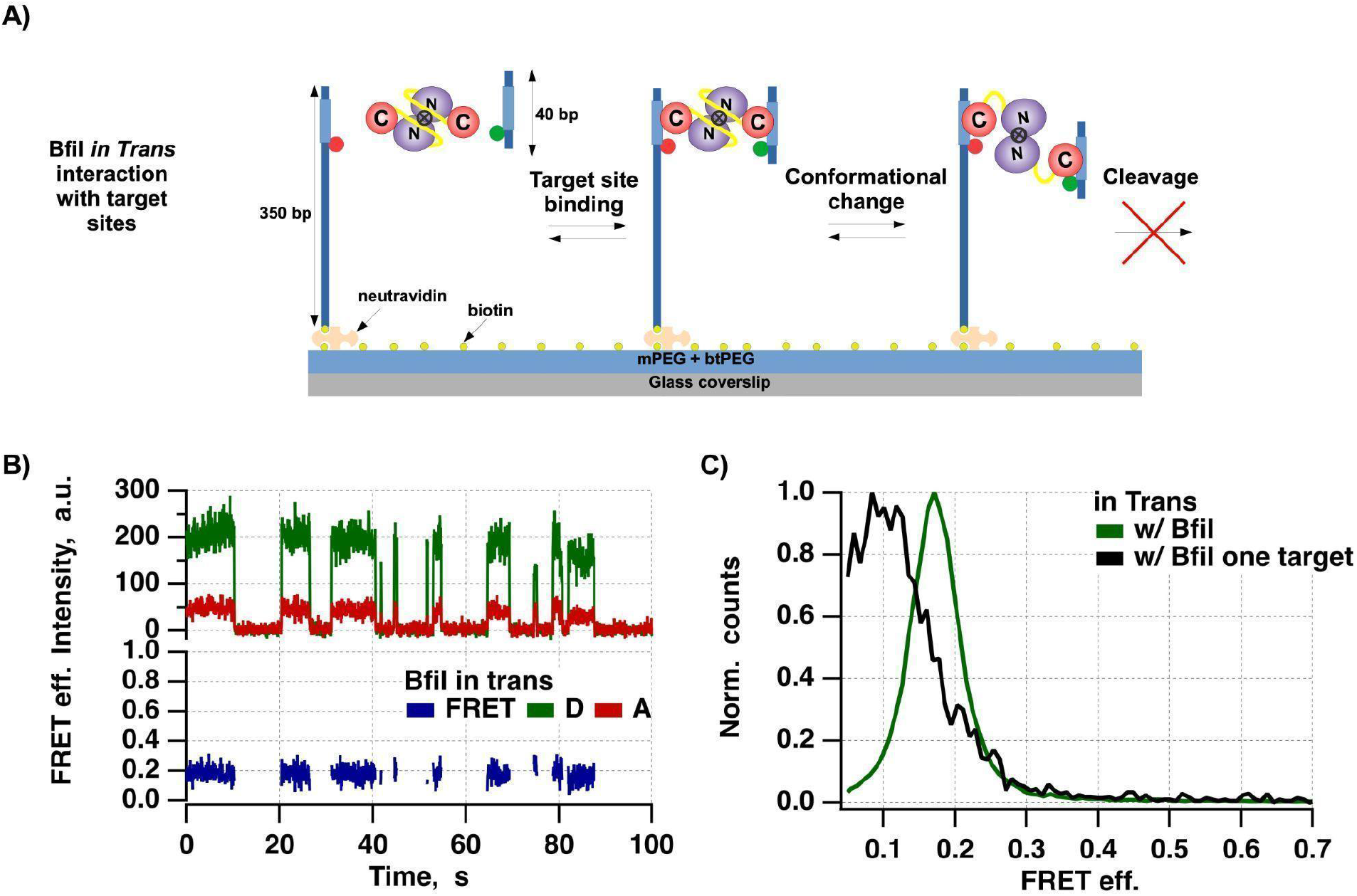
Results of single-molecule FRET (smFRET) BfiI-DNA in Trans interaction studies. **A)** Diagram illustrating the smFRET assay for BfiI-DNA in Trans interaction. Biotinylated double-stranded DNA (bt-dsDNA) fragments containing BfiI target site and FRET acceptor (ATTO647N) were immobilized on PEGylated (10% biotin-PEG) surface via neutravidin. The BfiI and FRET donor-labeled dsDNA oligonucleotide (Cy3B-oligo) were present in the solution. BfiI can either first bind the Cy3B-oligo or the immobilized bt-dsDNA. Binding of this complex to the non-target location of the bt-dsDNA results in “zero” FRET efficiency and high Cy3B fluorescence emission intensity, whereas binding to the target site results in increased FRET efficiency and lower Cy3B fluorescence emission intensity. Once BfiI is bound to the target, it potentially can change conformation from closed to open state. Such transition should change FRET efficiency from high to low. **B)** Representative SM signals: donor intensity (green), acceptor intensity (red), and apparent FRET efficiency (blue), showing DNA-BfiI interaction in Trans. **C)** Graph showing the distributions of FRET efficiencies of the detected binding events.

We first immobilized the biotinylated dsDNA molecules on a PEGylated (btPEG/mPEG) glass coverslip surface in a flow cell channel via neutravidin (SI Fig. 2A) and imaged them using the TIRF microscope with 532 and 635 nm wavelength excitations separately. Subsequent signal analysis resulted in a mean “zero” FRET level of ∼ 0.1 FRET efficiency.

We then bleached Cy3B labels on the DNA molecules (532 nm laser for ∼3 min) and acquired images with 532 and 635 nm excitations separately (SI Fig. 2B). We observed complete Cy3B bleaching and no evidence of ATTO647N bleaching, which was expected given the stability of ATTO647N dye in an oxygenated environment and its low absorption at 532 nm excitation. Next, we injected a mixture of BfiI preincubated with the Cy3B-oligo, together with the Tx and OS system and performed imaging as before (SI Fig. 2C). The BfiI:Cy3B-oligo ratio of 2.5:1 was chosen to maximize the fraction of BfiI molecules interacting with a single Cy3B-oligonucleotide, leaving the second DNA binding domain free to interact with the surface-immobilized DNA. Such an experimental scheme allowed us to perform long-lasting imaging and circumvent the donor bleaching because each different BfiI binding event is with a new donor fluorophore and the acceptor is not excited directly (SI Fig. 3).

The extracted FRET donor and acceptor SM signals showed fluorescence bursts of various duration that had clearly higher than noise Cy3B fluorescence emission intensities and FRET efficiencies (Fig. 4B). Examination of traces revealed that ∼20% of all signals represented BfiI binding events. To characterize these events, we performed semi-automated detection using criteria described in the materials and methods section. Distributions of FRET efficiencies that were made from all points of all detected binding events showed a distinct peak centered at ∼0.18 FRET efficiency (Fig. 4C).

To validate our findings, we used Cy3B-labeled dsDNA oligonucleotide without BfiI target site in otherwise identical experiments (Fig. 4C, SI Fig. 4). These experiments showed a 10-fold lower number of binding events, with their peak FRET efficiency centered at < 0.15. This negative control confirmed that on-target binding events must involve two BfiI recognition sites, and that off-target binding events are rare. Thus, we could easily isolate signal parts that reflected BfiI binding to the immobilized DNA and discriminate between off- and on-target site binding events even though they were closely spaced in the apparent FRET efficiency space.

### BfiI-DNA conformational dynamics

After we determined that BfiI was able to bind the target site *in Trans* on the immobilized DNA, we aimed to characterize conformational dynamics of BfiI upon DNA binding. We hypothesized that BfiI conformational transitions from “closed” to “open” state would result in a shift from high to low or “zero” FRET efficiency. To determine the occurrence of these conformational changes, we visually inspected each of the previously detected binding events (Fig. 5A-B). We attributed the binding events that did not exhibit any FRET efficiency changes to proteins that were not changing conformation during the binding event, and it could be either in closed or opened conformation (Fig. 5A).

**Figure 5:**
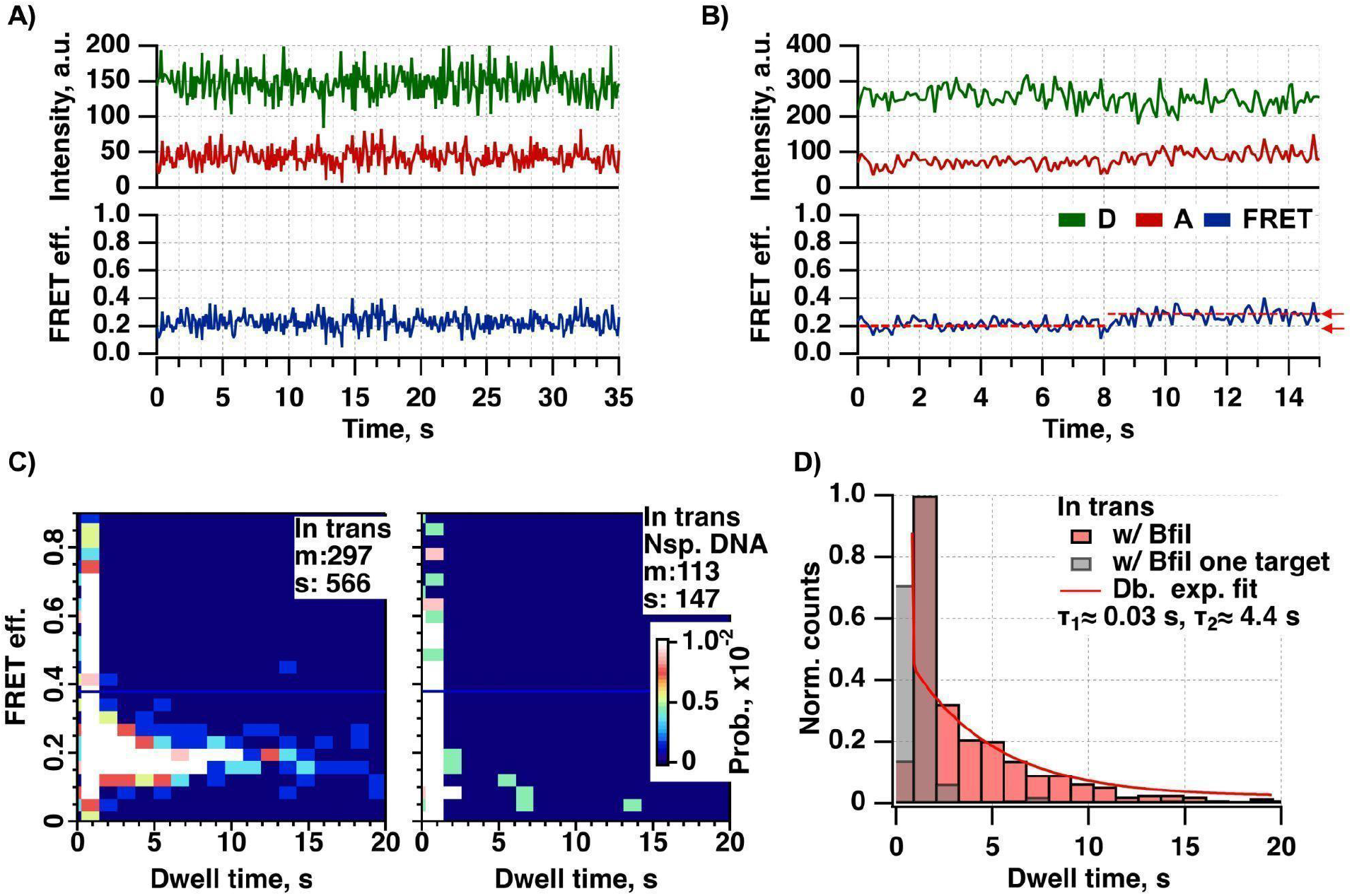
Conformational dynamics of BfiI occurring within binding events and binding events characterization. **A-B)** Graphs show single-molecule (SM) signals of donor (green) and acceptor (red) fluorescence emission intensities and FRET efficiencies (blue) of BfiI binding events with **A)** stable FRET efficiency, and **B)** step-wise changing FRET efficiency marked by red dashed lines and indicated by red arrows. **C)** 2D histogram plot correlating FRET efficiency of binding event with its duration for conditions w/ BfiI-oligoCy3B containing target site, and w/ BfiI-oligoCy3B and only a single target site. **D)** Horizontal line profiles for conditions w/ BfiI-oligoCy3B containing target site (red), and w/ BfiI-oligoCy3B and only a single target site (gray) taken over the corresponding 2D histogram plots shown in the panel **C**. The red-colored line-profile was fitted using a double-exponential function.

Based on the analysis of observed BfiI-DNA binding events, we calculated that approximately only 5% of all binding events were indicative of protein conformational changes (Fig. 5B). This suggested that (in relevant experimental conditions) BfiI conformational changes were not detectable when bound to the target.

Stable FRET efficiency within a binding event allowed us to reliably calculate the mean FRET efficiency of a binding event, which would be less meaningful in case of conformational changes within the binding event. To gain better understanding of binding events, we decided to correlate average FRET efficiency (i.e. BfiI conformation) and duration (stability of the complex) of the binding event. For this purpose we generated 2D histogram plots, where x-axis represented duration and y-axis – average FRET efficiency of the event (Fig. 5C). The plots visually demonstrated that in the presence of two target sites (both on the oligo DNA and immobilized DNA) binding events lasted longer than in the presence of a single target site (on immobilized DNA molecule, but not on the oligo in solution).

Double exponential function fitting to the binding events duration distribution showed a good fit (Fig. 5D). The short component lasted for 0.03 s, and the long component lasted for 4.4 s. In comparison to BfiI binding events with two target sites, the observed binding events were notably shorter when only one target site was present, and averaged at <0.5 s. The short binding events had variable FRET efficiencies, since the durations were too short to reliably determine their average FRET efficiency. Such binding events were also observed with the non-specific DNA substrate, and therefore they likely represent semi-specific protein interactions with DNA molecules. The apparent long duration (characteristic dwell time 4.4 s) events represented BfiI protein bound to two cognate target sites.

## Conclusion

Here we demonstrated application of two smFRET-based assays for studying BfiI-DNA interactions. The smFRET-based DNA looping assay, which we developed previously, was able to detect both “Phi”- and “U”-shaped DNA looping events as well as the non-looped DNA events *in Cis* for Ecl18kI and NgoMIV REases. However, this approach was not able to discriminate between different types of BfiI protein binding events occurring on an immobilized DNA molecule. Also, we could not not entirely rule out the possibility of different DNA loop conformations, which could potentially lead to different apparent FRET efficiencies of the looped DNA state. Besides, since both donor and acceptor fluorophores are always present, this smFRET-based DNA looping assay highly limits the meaningful observation time due to donor bleaching. In order to improve the encountered problems, we modified the previous assay to allow only *in Trans* BfiI-target DNA interactions. This assay employed FRET acceptor-labeled surface immobilized DNA and FRET donor-labeled dsDNA oligonucleotide, each containing a single binding target site. Dividing the BfiI target across two separate dsDNA molecules eliminated the possibility of DNA looping. This also allowed us to monitor the BfiI-target DNA interactions for longer durations, since every new BfiI binding to immobilized DNA event most likely happened with a new donor molecule.

Modified smFRET-based assay and TIRF microscopy allowed us to directly observe the BfiI-DNA binding events occurring on- and off-target. Our results showed that off-target binding events were short and lasted for < 0.5 s, while the on-target events were longer and lasted for > 4 s. We expected to detect BfiI conformational transitions from “closed” to “open” state upon cognate DNA binding by observing fluctuations of FRET efficiency from high to low. However, apparent FRET efficiency changes were rare, therefore we concluded several possible outcomes: 1) the BfiI protein was not able to perform conformational transitions under our experimental conditions; 2) the conformational change from “closed” to “open” state did not increase the distance between donor and acceptor fluorophores to meaningfully influence FRET signal; 3) the conformational transitions were happening too rapidly and could not be detected.

The modified assay we describe here in a way is similar to protein assisted-PAINT experiments,^41^ used for single stranded DNA labeling in super-resolution microscopy studies, whereas BfiI could be utilized for dynamic labeling of dsDNA substrates, in “BfiI-PAINT’’ experiments. While the Argonaute protein can be loaded with a guide molecule of desired sequence (thus allowing more controlled labeling), the dsDNA substrate should contain at least one BfiI target sequence for successful labeling. We believe that this dsDNA labeling approach could prove useful for DNA Curtains experiments ^38,42–45^ and other DNA stretch-assays.^11,46^ Besides dsDNA labeling, it is likely that our developed assay will be useful for other two targets-binding DNA-interacting proteins.

## ASSOCIATED CONTENT

## Funding Sources

This project has received funding from the Lithuanian Research Council (S-MIP-20-55 for M.T.).

## ACKNOWLEDGMENT

We are grateful to Prof. Dr. Virginijus Šikšnys for valuable advice and discussion throughout this work.

## Supporting Information

The Supporting Information is available free of charge.

## Notes

### Competing Interest Statement

The authors have declared no competing interest.

