## Supplemental info file for "Advancing Understanding of DNA-BfiI Restriction Endonuclease Cis and Trans Interactions through smFRET Technology"

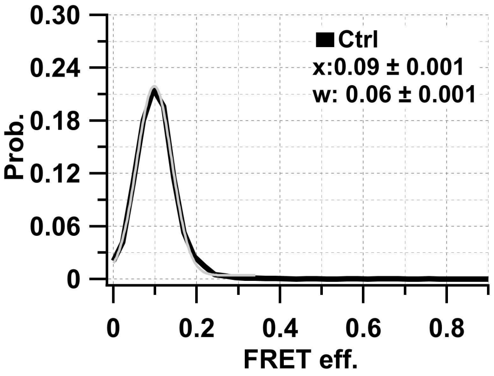


**SI Figure 1.** FRET efficiency of Cy3B and ATTO647N labeled biotinylated double-stranded DNA immobilized on PEGylated surface via neutravidin before bleaching step.

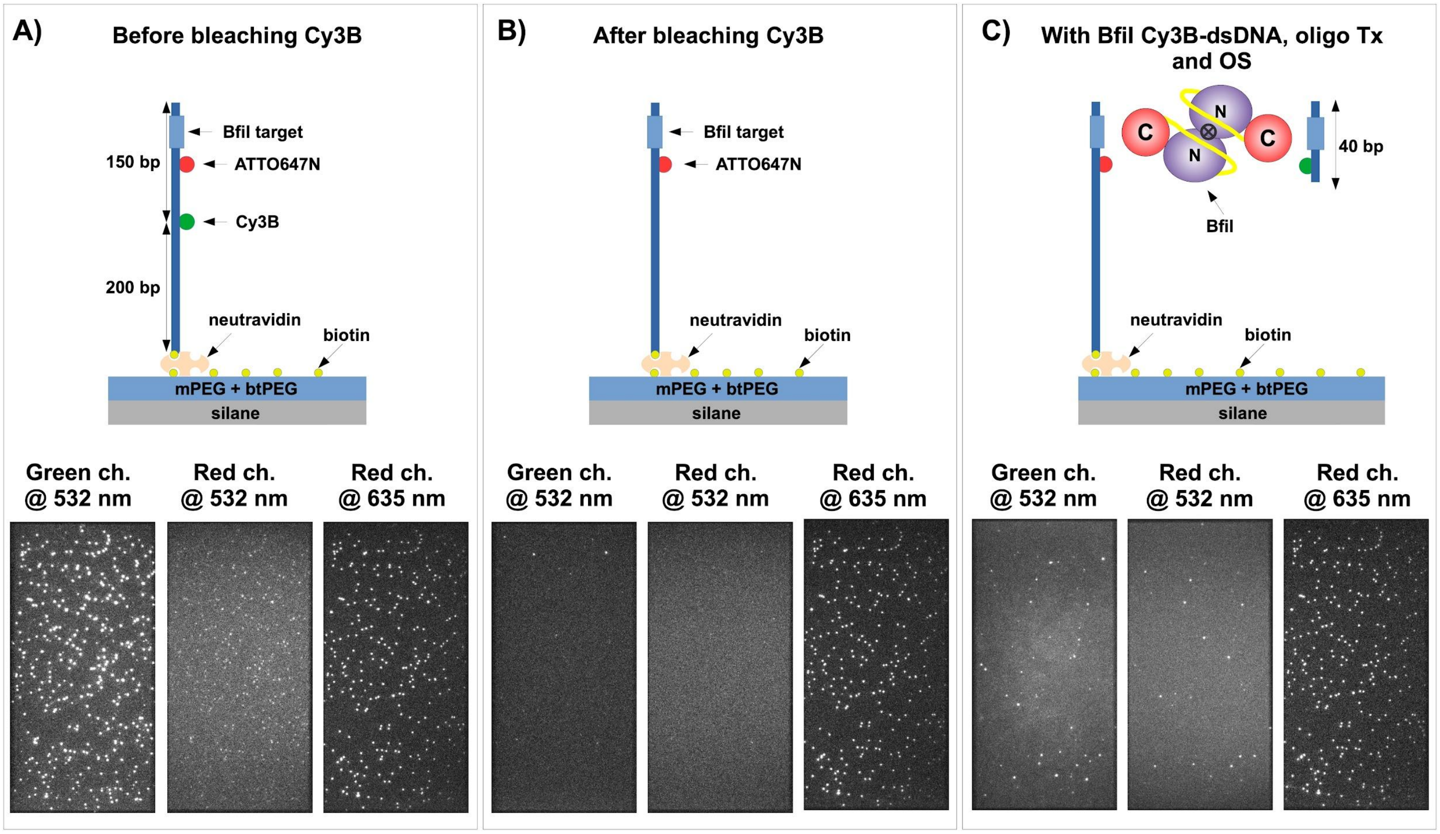


**SI Figure 2:** Diagrams illustrating the scheme of single-molecule FRET (smFRET) assay for BfiI-DNA in Trans interaction. **A)** Cy3B and ATTO647N labeled biotinylated dsDNA containing a single BfiI target site was immobilized on a PEGylated glass coverslip surface via neutravidin. The TIRF images acquired under 532 and 635 nm excitations showed co-localizing fluorescent spots. **B)** Cy3B on the biotinylated dsDNA was bleached out by shining 532 nm laser for 3 min. The TIRF images acquired under 532 and 635 nm excitations showed completely bleached Cy3B and no evidence of ATTO647N bleaching. **C)** Binding of BfiI was monitored by injection of BfiI and Cy3B-labeled double-stranded DNA oligonucleotide (Cy3B-oligo) in presence of Trolox (Tx) and oxygen scavenger (OS) system. The TIRF images acquired under 532 and 635 nm excitations showed increased background of Cy3B fluorescence and some fraction of co-localizing Cy3B and ATTO647N fluorescent spots.

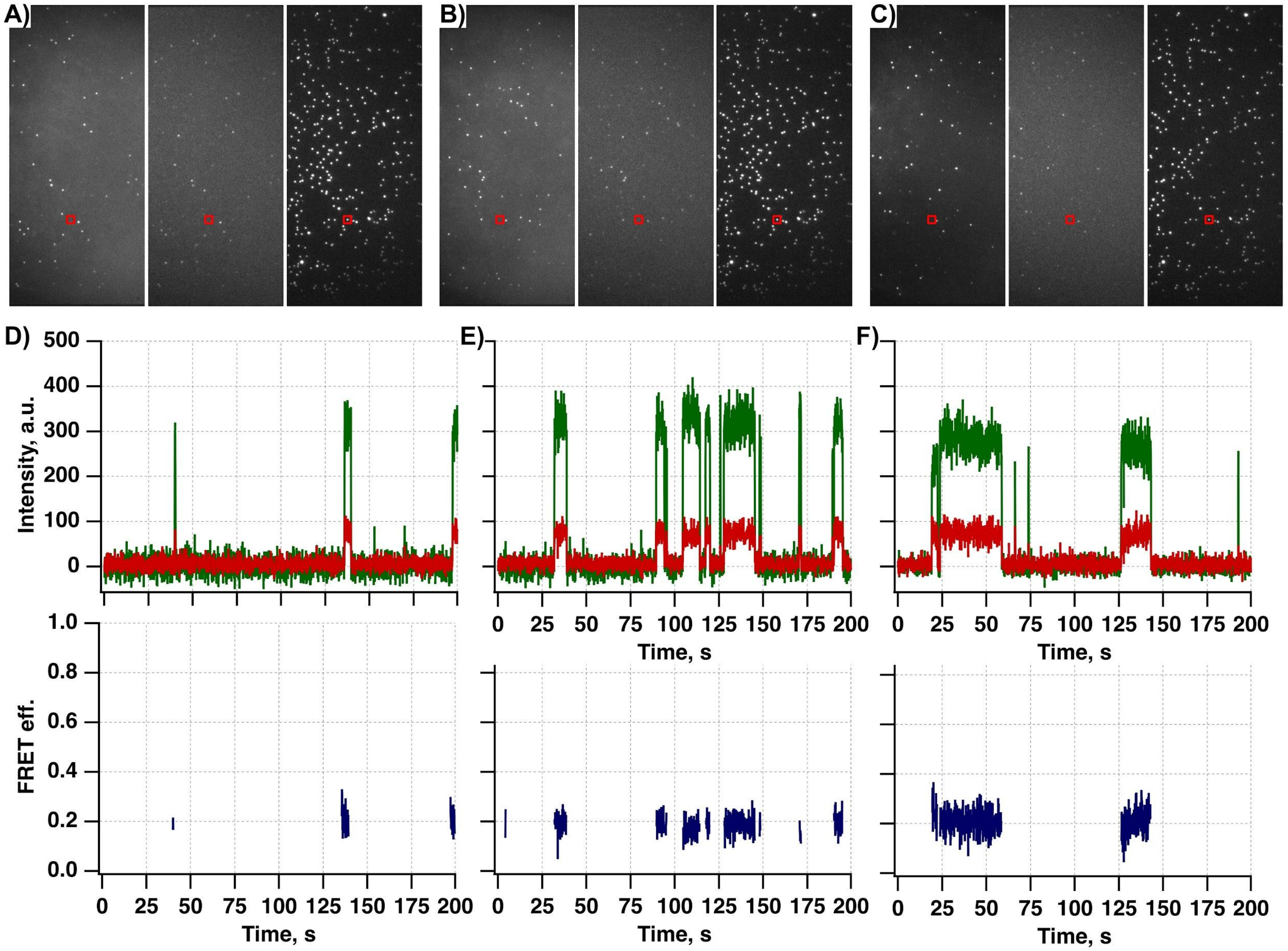


**SI Figure 3.** The long-lasting imaging of DNA-BfiI interaction In trans. 0.5 nM of BfiI and 0.2 nM of Cy3B-labeled dsDNA oligonucleotide with BfiI target site together with Tx and OS system incubated with the surface-immobilized single BfiI target containing DNA molecules. **A)** Illustrative single-molecule donor (left micrograph) and acceptor (middle micrograph) images that were acquired at 532 nm wavelength excitation and acceptor (right micrograph) image that was acquired at 635 nm wavelength excitation. **B)** The same surface position as in panel **A**, just after 200 s. **C)** The same surface position as in panel **B**, just after 200 s. **D-F)** Illustrative single-molecule donor (green) and acceptor (red) fluorescence intensity traces that were acquired under described conditions at 532 nm wavelength excitation along with the apparent FRET efficiency (blue) traces. These traces were extracted from the fluorescent spot indicated by red squares in panels **A**, **B**, **C**. Panel **D** corresponds to panel **A**, panel **E** corresponds to panel **B**, and panel **F** corresponds to panel **C**.

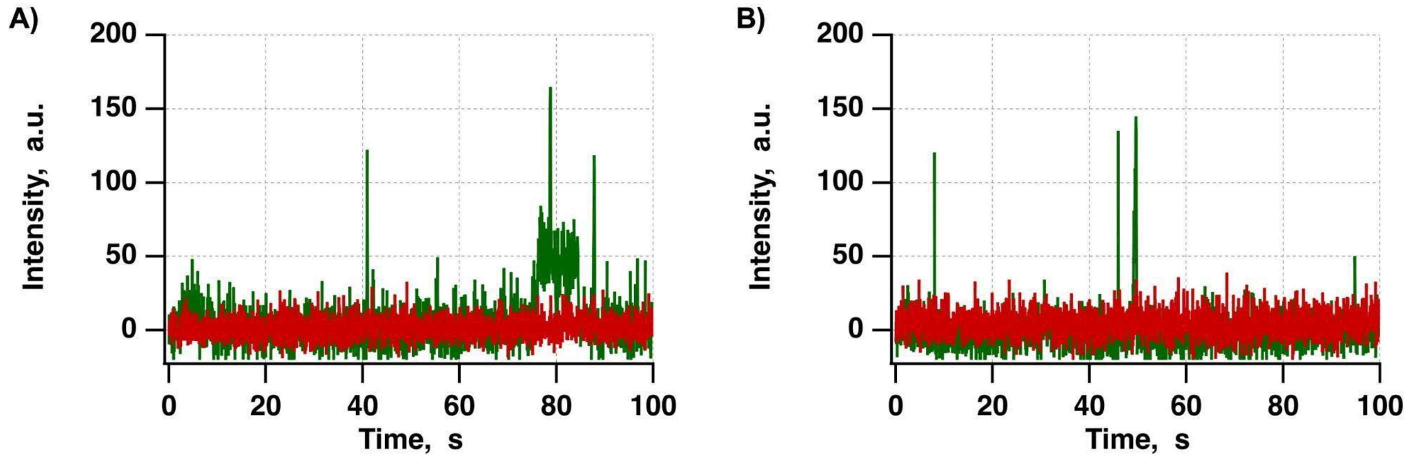


**SI Figure 4.** DNA-BfiI interaction In trans negative control. 0.5 nM of BfiI and 0.2 nM of Cy3B-labeled dsDNA oligonucleotide without BfiI target site together with Tx and OS system incubated with the surface-immobilized single BfiI target containing DNA molecules. A-B) Illustrative single-molecule donor (green) and acceptor (red) fluorescence intensity traces that were acquired under described conditions at 532 nm wavelength excitation.
